# Synonymous and single nucleotide changes facilitate the adaptation of a horizontally transferred gene

**DOI:** 10.1101/2024.09.20.614038

**Authors:** Pavithra Venkataraman, Sudeepta Singh, Neetika Ahlawat, Supreet Saini

## Abstract

The movement of genes between microbial species, or horizontal gene transfer, is common. While this process speeds up adaptation, the functionality of horizontally transferred genes is highly constrained in the new hosts due to several reasons, and protein localization is one of them. In this study, we ask what is the minimum number of mutations that can resolve a localization problem faced by a horizontally transferred protein in a new host. Using a directed-evolution approach, we show that SNPs and a synonymous mutation can change the localization patterns of an ammonia transporter (AmtA) moved from *Dictyostelium discoideum* to *Saccharomyces cerevisiae*. Interestingly, the mutations that cause this change in localization, confer different fitness effects, are spread throughout the gene, and do not cause a uniform change in amino acids. In a novel attempt, we show how SNPs or a synonymous mutation can alter the membrane affinity of proteins, and as a result, aiding protein evolution and functional diversification.

## Background

Horizontal gene transfer (HGT) is ubiquitous and is an important mode by which new phenotypes are acquired in living organisms^1,2^. Adapting via this mode is “quick”, relative to heritable genetic changes, and thus HGT speeds up evolution^3^. HGT is also relevant in the context of functional diversification - in a new host, the horizontally transferred gene can confer a new trait by itself, or by undergoing genetic changes. Beneficial horizontally transferred genes will be retained by the new host, while deleterious ones will be purged by selection^3^.

The function of the transferred gene, the protein’s interaction network, the differences in the cellular environment of the donor and recipient, and interactions of the foreign gene with the host genome are thought to pose potential barriers to the success of horizontal gene transfer^4^. A combination of these factors mostly leads to the loss of the horizontally transferred gene in the new host, either due to selection or drift. However, ubiquitous mechanisms of gene silencing exist, indicating that neutral or deleterious genes could be retained in the cell until they adapt^5^. In such a scenario, if multiple short adaptive routes are available for a gene to gain functionality, there are greater chances that a foreign gene will be retained. Conversely, if many mutations are needed before the gene gains functionality, its evolutionary fate is largely drift-dependent. In this context, what is the number of mutations that would make a non-functional horizontally transferred gene functional?

To answer this question, in this work, we mimic an HGT event by shifting an ammonia transporter from *Dictyostelium discoideum* to a *Saccharomyces cerevisiae* strain lacking ammonia transporters, and ask how “easy” it is, in terms of the nature and number of mutations, for the horizontally transferred gene to gain functionality in a new host.

Ammonia transporter proteins (Amt/MEP/Rh) are present across all life forms and are believed to have a common origin^6^. Most bacteria and fungi have Amt or MEP proteins, while most mammals have Rh proteins. Depending on the organism and environment, these membrane proteins have been shown to transport ammonia, methylamine, and carbon dioxide^7^. In the yeast *Saccharomyces cerevisiae*, there are three MEP proteins (MEP123), which localize in the plasma membrane^8^, and transport ammonia into the cell. One of them, MEP2, also serves as an ammonia sensor and mediates pseudohyphal differentiation^9^.

In strains lacking these three MEP proteins (triple-mutants), successful functional complementation of ammonia transporters from other organisms like *Escherichia coli* and *C. elegans*, via horizontal gene transfer, has been reported^10–13^. However, an ammonia transporter from *Dictyostelium discoideum*, AmtA, cannot functionally complement the triple-mutant *S. cerevisiae* cells^14^. Subcellular localization of the Amt proteins (AmtA, AmtB, and AmtC) in *D. discoideum* showed that AmtA and AmtB localize on the membranes of endolysosomes and phagosomes, and export ammonia^15^. It has been shown that AmtA suffers from a localization problem when transferred to the triple-mutant yeast strains and that a combination of mutations is necessary for it to be functional in the new host^14^.

The goal of this study is to identify the least number of genetic changes that are required for the non-functional horizontally transferred gene to become functional in the new host. In the context of AmtA from *Dictyostelium*, we ask what is the nature and number of mutations required for it to import extracellular ammonia into a yeast cell. Through this work, we aim to provide insights on the evolutionary history of ammonia transporters by asking how “easy” it has been for these genes to shift hosts and be functional. In this context, we isolate functional mutants of AmtA that have the minimum number of mutations (one SNP), via a directed-evolution approach, and characterize their fitness effects in yeast.

## Results

### AmtA suffers from a localization problem in *S. cerevisiae*

Triple mutant *S. cerevisiae* cells with AmtA (driven by MEP2-promoter) did not form CFUs in low ammonia (500uM) environments (**Figure 1**). This could be due to one of two reasons: one, AmtA protein is unstable in yeast, or AmtA does not localize in the yeast cell membrane. To investigate which of the two possibilities makes AmtA non-functional in yeast, we transformed the triple mutant yeast cells with an AmtA-GFP construct (driven by MEP2-promoter). We grew these cells in low ammonia conditions. Upon performing confocal microscopy, we observed that these proteins localized on the membranes of cell organelles like they did in *D. discoideum* (**Figure 2**). Clearly, MEP2 and AmtA localize differently in yeast.

**Figure 1.**
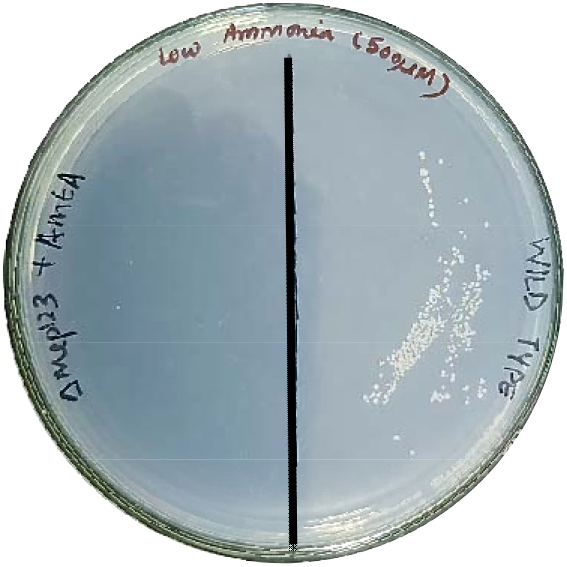
AMTA does not functionally complement *S. cerevisiae* lacking MEP123. (Left) MEP123Δ complemented with AmtA is not able to form colonies on a plate with low ammonia as the sole carbon source. (Right) Ancestor strain is able to form colonies on media containing 500μM Ammonia as the sole nitrogen source.

**Figure 2.**
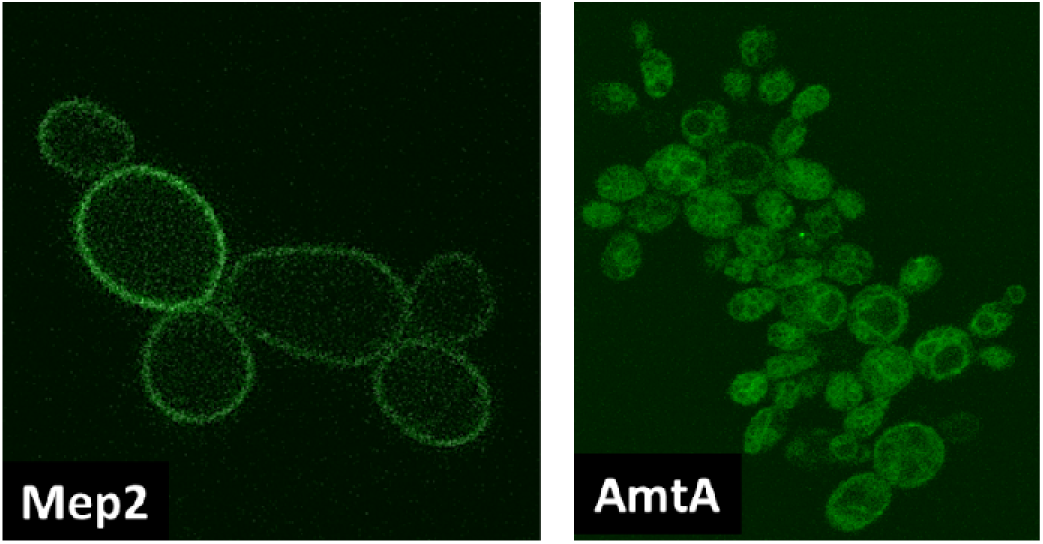
MEP2 (left) localizes on the cell membrane. AmtA (right) localizes intracellularly on membranes of organelles and in the cytoplasm.

### Single non-synonymous mutations in AmtA alter its localization patterns in *S. cerevisiae*, and their fitness effects are environment-dependent

Using a directed-evolution approach (Methods), we screened about five hundred AmtA alleles and isolated **nine** functional variants. The mutants were isolated based on their ability to form colonies on low ammonia plates (**Figure 3**). The functional AmtA mutants that grew on low-ammonia (500uM) plates upon streaking took between 4 and 8 days to form single colonies (see Methods). Therefore, it was clear that the fitness conferred by these mutations are different.

**Figure 3.**
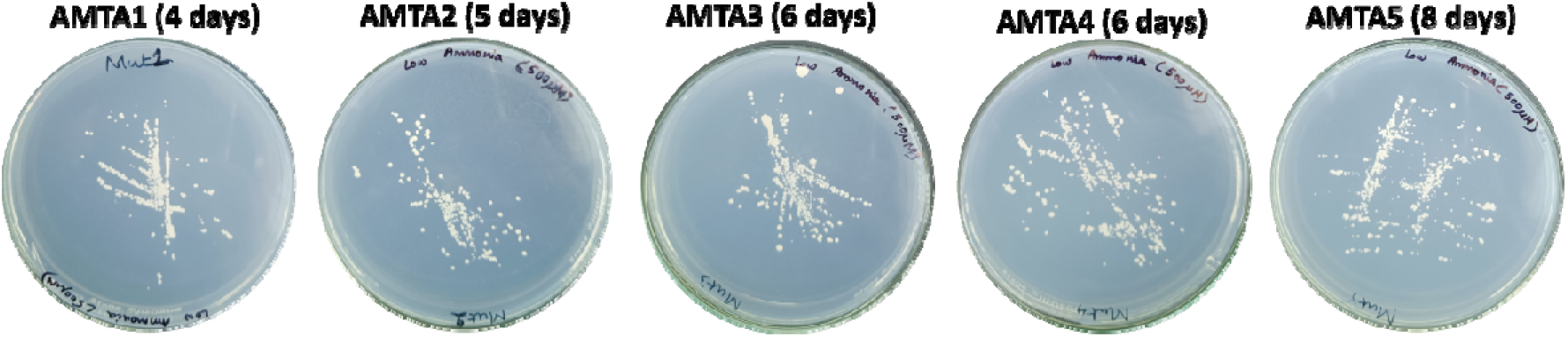
Functional AmtA mutants in yeast form colonies on a low ammonia plate. The five AmtA mutants carrying one mutation each take different amounts of time to form colonies on 500uM ammonia plates.

We Sanger sequenced the functional AmtA alleles to identify the nature and number of mutations. As shown in **Table 1**, five of the nine alleles have only distinct mutation in each (AmtA1-AmtA5), and one of them is a synonymous mutation (AmtA5). We isolated two triple mutants (AmtA6, AmtA7) and two double mutants (AmtA8, AmtA9). These mutations are not localized to any region, or do not represent a particular type of change.

**Table 1.** Mutant library screen led to isolation of nine functional variants of AmtA in yeast. Five out of the nine variants carried only a single SNP. One of the five variants carried only a synonymous mutation. The nature of change of amino acid or the location of the mutations follow no specific trends. The batch culture growth rates of the yeast strains carrying these functional AmtA alleles are environment-dependent.

| Allele | Amino acid change | R group change | Growth rate, relative to WT (500uM) | Growth rate, relative to WT (5uM) |
| --- | --- | --- | --- | --- |
| AMTA1 | P55H | Polar uncharged to positively charged | $1.17 \pm 0.045$ | $1.60 \pm 0.14$ |
| AMTA2 | S350G | Polar uncharged to non-polar aliphatic | $0.62 \pm 0.053$ | $1.04 \pm 0.21$ |
| AMTA3 | L304S | Non-polar to polar uncharged | $0.64 \pm 0.038$ | $0.72 \pm 0.06$ |
| AMTA4 | F97L | Aromatic to non-polar aliphatic | $0.45 \pm 0.0084$ | $0.68 \pm 0.091$ |
| AMTA5 | L284L | NA | $0.37 \pm 0.0021$ | $0.55 \pm 0.034$ |
| AMTA6 | V43D, S67S, V86A | Non-polar aliphatic to Negatively charged, NA, Non-polar aliphatic to non-polar aliphatic | $1.24 \pm 0.08$ | $1.49 \pm 0.039$ |
| AMTA7 | P55L, N315S, A329T | Polar uncharged to non-polar aliphatic, Polar uncharged to polar uncharged, Non-polar aliphatic to polar uncharged | $1.74 \pm 0.125$ | $2.64 \pm 0.28$ |
| AMTA8 | P231S, L284L | Polar uncharged to polar uncharged, NA | $0.35 \pm 0.022$ | $0.49 \pm 0.023$ |
| AMTA9 | W88R, I364T | Aromatic to positively charged, nonpolar aliphatic to polar uncharged | $0.98 \pm 0.075$ | $1.08 \pm 0.039$ |

Since the focus of this work are the single mutants, we fused GFP at the C-terminal ends of single-mutant functional AmtA alleles, and transformed triple mutant yeast cells with the construct. Upon performing confocal microscopy, we observed that only the plasma membrane fluoresced, as shown in **Figure 4**, indicating that there are several mutations that solve the localization problem that the wild-type AmtA was suffering from in yeast.

**Figure 4.**
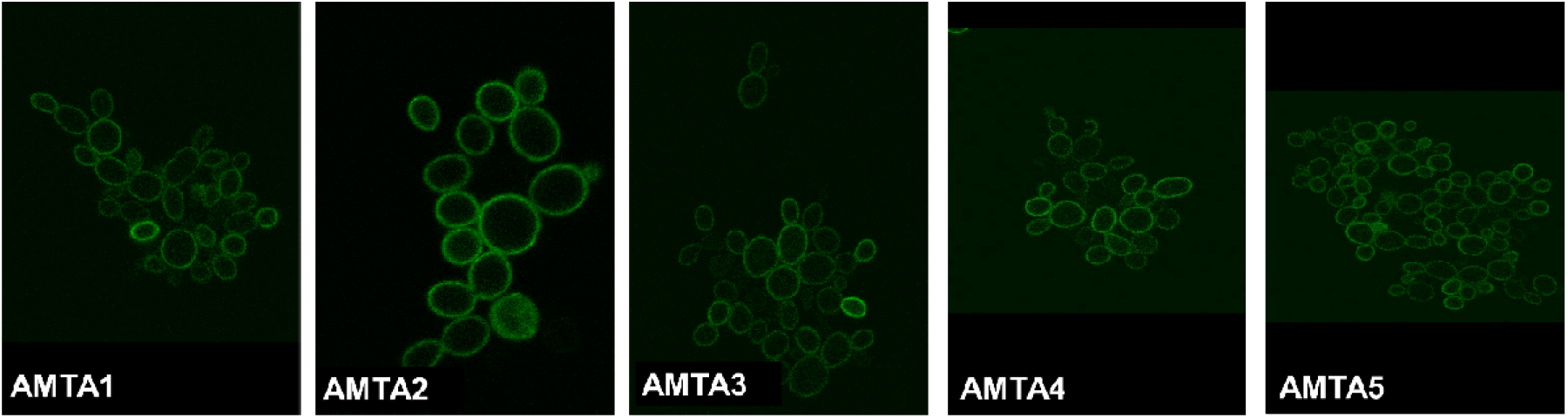
A single SNP changes the pattern of localization of AmtA in yeast. Confocal microscopy shows that acquisition of one SNP in AmtA sequence changes the protein’s localization from cell organelle membrane to the plasma membrane. AmtA5 achieves this via the acquisition of a single synonymous mutation.

To identify the fitness effects of the functional AmtA alleles and their dependence on environment conditions, we measured the batch culture growth rates (see Methods) of the triple-mutant yeast strain carrying the functional alleles in two concentrations of ammonia (500uM & 5uM). As shown in **Table 1**, the fitness effects of these beneficial mutations form a spectrum, and are significantly different from each other. Although these functional alleles were isolated based on growth in 500uM ammonia plates, their fitness effects relative to the wild-type was higher in 5uM ammonia environment than 500uM. This indicates that along with the host environment, the external environment in which the host is present influences the likelihood of retention of the foreign gene.

## Discussion

Horizontal gene transfer has been implicated in a range of studies involving the spread of antibiotic resistance, virulence, and the evolution of new traits^16^. The success of HGT is contingent on whether it gets inserted at an appropriate position in the host genome and if it gets transcribed, translated, and trafficked accurately^17^. In several cases, genes enter new hosts and confer an immediate adaptive benefit, and their retention in the new host is favored by selection. However, in cases where there is no immediate adaptive benefit, the success of a horizontal gene transfer event is unpredictable^4^.

One way that a non-functional gene can manage to get retained in the new host is by adapting “quickly”, and conferring a fitness benefit. In this work, we ask if such options exist by horizontally transferring a *D. discoideum* ammonia transporter protein (AmtA) to *S. cerevisiae* strains that lack their native ammonia transporters (MEP1, MEP2, MEP3). The foreign gene is driven by the native yeast MEP2 promoter, so our results are not biased by regulatory constraints^17^.

Our results show that there are several adaptive SNPs, including a synonymous mutation, that are available to AmtA in the yeast *S. cerevisiae*. While the fitness effect of these mutations is diverse, they all appear to flip the membrane curvature affinity (convex or concave) of the protein, and thus solve the localization problem. We found only one similar observation in literature, where a mutation associated with interstitial lung cancer alters the subcellular localization of a membrane protein in humans^18^. When taken together with recent evidence from laboratory experiments, our results highlight that synonymous mutations have context-dependent effects on fitness^19–23^. The implications of change in the localization patterns of AmtA via SNPs, in this context, are on (a) protein evolution, (b) directionality of protein function, and (c) success of HGT. The evolvability of functional AmtA variants in yeast remains to be tested.

We observe that the fitness effects of our functional AmtA alleles are not uniform. While a naive hypothesis would be that these mutations confer different translation and localization dynamics, in the absence of any repeatability in changes of amino acids (see Figure S1), we are unable to comment on the exact mechanism driving these differences in fitness effects. In fact, it is also unknown how these SNPs alter the localization pattern of the protein in the new host because the mutations are not localized to any region, including the signal peptide^24,25^. Therefore, our work motivates further research into correlating genetic changes with protein localization mechanisms.

It has been hypothesized in the past that operational genes are more likely to be successfully horizontally transferred than informational genes^26^. Our results support this hypothesis and provide quantitative insights into the number of mutations that would make a standalone gene functional in a new host. Broadly, our results also explain how a set of orthologous proteins could have evolved to perform a diverse range of functions over evolutionary timescale by shifting hosts.

## Materials and methods

### Strain and plasmid used

MLY61**a** was used as the ancestor, and MLY131**a** was used as the MEP123 triple deletion strain^9^. AmtA was cloned between the SalI and BamHI sites and MEP2 promoter was cloned between HindIII and SalI cites in the plasmid YCPlac33. GFP tagged AmtA was cloned as previously described^14^.

### Media conditions used

Low ammonia plates were made using the following recipe: 2g of glucose, 0.17g of DIFCO Yeast Nitrogen Base without amino acids and ammonium sulphate, 2g of agar and 1mL of 50mM ammonium sulphate were added to 100mL of Milli-Q water and sterilized by autoclaving.

### CFU formation on low ammonia plates

Cells were streaked on a uracil dropout media (2% glucose, 0.05% ura dropout media, 0.66% YNB containing Ammonium Sulphate), and incubated at 30 deg C. The colonies on the plate were then streaked on media with low ammonia as the sole nitrogen source (2% glucose, 0.17% YNB without ammonium sulphate and amino acids, and ammonium sulphate to a final concentration of 500μM). The plates were then incubated at 30 deg C and appearance of colonies noted.

### Microscopy

Cells were grown in low ammonia culture media for ∼40 hours. The supernatant was removed by spinning the down the cells at 3000g for 3-4 minutes. The cell pellet was washed with 0.5 ml PBS buffer. The cells were spun again and resuspended in 100μl PBS. A volume of 5μl was spread on a glass slide, and covered with a cover slip. The cells were then imaged at 63x/1.4NA (oil) in a Laser Scanning Microscope (Carl Zeiss, LSM780).

### Library creation and mutant selection

Error-prone PCR was used to generate mutant AMTA genes, which were later screened for functionality (the ability to form colony on a low ammonia plate). The following recipe was used for a 50μL error-prone PCR mix: Forward primer (30μM) - 1μL; Reverse primer (30μM) - 1μL; Magnesium chloride - 10μL; dNTPs - 0.5μL; Thermo Scientific Taq Green Master mix (2X) - 25μL; Template - 0.5μL; Manganese chloride - 2.5μL, 5μL and 7.5μL; Distilled water - as required to make up reaction volume to 50μL.

The PCR product was purified and mixed with the plasmid pRI51 cut with BamHI and SalI. The mix was transformed in MEP123Δ strain, and played on ura-plates for individual colonies. The colonies were streaked on plates with 500μM ammonia and screened for the variants which form colonies.

### Quantification of fitness

Each of the functional mutants was streaked on a ura-plate. After a colony formed, a small part of the colony was used to inoculate 5mL liquid ura-liquid media. This culture was allowed to grow at 30ºC until an O.D(600nm) of about 1 is reached. Then, 10mL of low ammonia (500μM) medium is inoculated with the primary culture, such that the O.D(600nm) was 0.01. This culture was allowed to grow at 30ºC, and the O.D(600nm) measured every 4 hours, to obtain the growth curve. Wild-type *S. cerevisiae* cells with pRI51 was used as the positive control and triple mutant strain with pRI51 served as the negative control.

### Sequencing

AmtA fragments were PCR-amplified, and Sanger sequenced at Allied Scientific Products & Eurofins Scientific.

## Acknowledgements

We thank Sergey Kryazhimskiy for feedback on this work. We thank the International Centre for Theoretical Sciences for supporting the Bangalore Schools on Population Genetics and Evolution (code: ICTS/popgen2020/01) for facilitating discussions regarding this work.

